# The association of Plk1 with the Astrin-Kinastrin complex promotes formation and maintenance of a metaphase plate

**DOI:** 10.1101/2020.07.01.181933

**Authors:** Zoë Geraghty, Christina Barnard, Pelin Uluocak, Ulrike Gruneberg

## Abstract

Errors in mitotic chromosome segregation can lead to DNA damage and aneuploidy, both hallmarks of cancer. To achieve synchronous error-free segregation, mitotic chromosomes must align at the metaphase plate with stable amphitelic attachments to microtubules emanating from opposing spindle poles. The Astrin-Kinastrin/SKAP complex, also containing DYNLL1 and MYCBP, is a spindle and kinetochore protein complex with important roles in bipolar spindle formation, chromosome alignment and microtubule-kinetochore attachment. However, the molecular mechanisms by which Astrin-Kinastrin fulfils these diverse roles are not fully understood. Here we characterise a direct interaction between Astrin and the mitotic kinase Plk1. We identify the Plk1-binding site on Astrin as well as four Plk1 phosphorylation sites on Astrin. Regulation of Astrin-Kinastrin by Plk1 is dispensable for bipolar spindle formation and bulk chromosome congression but promotes stable microtubule-kinetochore attachments and metaphase plate maintenance. It is known that Plk1 activity is required for effective microtubule-kinetochore attachment formation, and we suggest that Astrin phosphorylation by Plk1 contributes to this process.

**Summary:** We demonstrate that Plk1 binds to and phosphorylates the N-terminus of Astrin. This interaction promotes recruitment of the Astrin-complex to kinetochores and stabilises microtubule-kinetochore-attachments in situations when mitosis is delayed.

## Introduction

During animal cell division one of the key prerequisites for successful chromosome segregation is the efficient end-on attachment of kinetochores to microtubules. This process is mediated both by proteins at the kinetochore and proteins delivered by the incoming microtubules, and is highly regulated by both kinases and phosphatases (Foley et al., 2011; Godek et al., 2015; Monda and Cheeseman, 2018). One of the key mitotic kinases required for the efficient formation and maintenance of kinetochore (K-) fibres is the Polo-like kinase 1 (Plk1) (Lenart et al., 2007). In mitosis, Plk1 is localised to centrosomes, centromeres and kinetochores, the spindle midzone and the midbody and has important phosphorylation targets at all of these sites (Arnaud et al., 1998; Barr et al., 2004). Localisation of Plk1 to specific loci during cell division is achieved by binding of the C-terminal polo-box-domain to phosphorylated docking sites containing a consensus S-[pS/pT]-P/X motif (Elia et al., 2003a; Elia et al., 2003b). Priming phosphorylations of PBD docking sites are often carried out by CDK1-cyclin B complexes but can also be generated by Plk1 itself (Elowe et al., 2007; Kang et al., 2006; Neef et al., 2007). At the kinetochore, proteins of the outer as well as the inner kinetochore have been described as binding partners for Plk1, creating specific local pools of Plk1 activity (Lera et al., 2016). How Plk1 supports the establishment of microtubule-kinetochore attachments is still not completely clear. One well described role of Plk1 at the kinetochore is to promote the binding of the PP2A-B56 phosphatase to the spindle assembly checkpoint protein BubR1 by phosphorylating the Kinetochore attachment Regulatory Domain (KARD) in BubR1 (Suijkerbuijk et al., 2012). BubR1-bound PP2A-B56 is then thought to help form stable microtubule-kinetochore attachments by opposing Aurora B activity (Foley et al., 2011; Suijkerbuijk et al., 2012; Xu et al., 2013). However, because of the multitude of known substrates and binding partners it is highly likely that there are other key substrates of Plk1 at the kinetochore, the functionality of which is not understood yet.

One such described interaction partner for Plk1 at the kinetochore is the mitotic spindle- and kinetochore-localised protein Astrin (Dunsch et al., 2011; Kettenbach et al., 2018). Astrin is a long coiled-coil protein with a globular N-terminal domain which forms a tetrameric complex with the kinetochore associated Astrin binding partner Kinastrin (also known as small kinetochore associated protein SKAP), dynein light chain 1 (DYNLL1) and the c-Myc binding protein MYCBP (Dunsch et al., 2011; Friese et al., 2016; Gruber et al., 2002; Kern et al., 2017; Schmidt et al., 2010). Depletion of Astrin or Kinastrin results in severe impairment of bipolar spindle formation, failure of chromosome congression and mitotic arrest (Dunsch et al., 2011; Gruber et al., 2002; Schmidt et al., 2010; Thein et al., 2007). In particular, the formation of stable, end-on microtubule-kinetochore attachments is impaired in Astrin depleted cells (Dunsch et al., 2011; Shrestha et al., 2017). The key microtubule binding protein complex at the outer kinetochore is the NDC80 complex (Cheeseman et al., 2006; Ciferri et al., 2008), and it has recently been reported that Astrin may help to stabilise attachments by binding synergistically with microtubules to NDC80 at the kinetochore (Kern et al., 2017). This role is promoted by Astrin delivering a specific pool of the PP1 phosphatase to kinetochores aiding with further Astrin enrichment (Conti et al., 2019). However, whether the Astrin complex cooperates with Plk1 in stabilising microtubule-kinetochore attachments has so far not been explored.

Here we characterise a direct interaction between Astrin and the mitotic kinase Plk1 and identify the Plk1-binding site on Astrin as well as four novel Plk1 phosphorylation sites in the N-terminus of Astrin. We find that regulation of Astrin by Plk1 is not required for most aspects of Astrin-Kinastrin complex functionality but does promote the maintenance of a stable metaphase plate. We hypothesise that the Astrin-Plk1 interaction safeguards faithful chromosome segregation in situations where mitotic progression is slowed.

## Results and Discussion

### Plk1 associates with the N-terminus of Astrin

Recent proteomic analyses of mitotic Astrin or Plk1 complexes, respectively, identified Plk1 as an interaction partner of Astrin (Dunsch et al., 2011; Kettenbach et al., 2018). To understand when and where during the cell cycle Astrin and Plk1 interact indirect immunofluorescence analysis of Astrin and Plk1 was employed. Untreated HeLa cells at prometaphase or metaphase or HeLa cells arrested in prometaphase-like states by treatment with the microtubule depolymerising drug nocodazole or the small molecule Eg5 kinesin inhibitor S-Trityl-L-Cysteine (STLC) (Skoufias et al., 2006) were stained for Astrin and Plk1. This analysis confirmed that Plk1 associated with both centrosomes and kinetochores in mitosis. Consistent with previous reports, Plk1 kinetochore staining was strongest in prometaphase and on unattached kinetochores and weaker at attached kinetochores at metaphase plates or in STLC-arrested cells (Lenart et al., 2007) (Figure 1A). In contrast, Astrin localised to spindle poles and decorated attached kinetochores, as previously reported (Mack and Compton, 2001; Manning et al., 2010; Schmidt et al., 2010; Thein et al., 2007). In metaphase cells or cells arrested with monopolar spindles as a consequence of treatment with STLC, Astrin and Plk1 co-localised at attached kinetochores (Figure 1A, inset panels at bottom). We therefore conclude that Astrin and Plk1 most likely interact at attached kinetochores as found physiologically at aligned metaphase plates or experimentally in monopolar, STLC arrested spindles, in which on average half of the kinetochores are attached. In cells depleted of Astrin, Plk1 was still visible at the kinetochores of the disorganised spindles (Figure 1B), in line with the idea that the bulk of kinetochore-localised Plk1 is bound by other proteins, such as BubR1 and Bub1 (Elowe et al., 2007; Jia et al., 2016; Wong and Fang, 2007). In contrast, Astrin was lost from kinetochores in Plk1 depleted cells (Figure 1B), most likely because in the absence of Plk1 no stably attached kinetochores can be generated (Lenart et al., 2007).

**Figure 1.**
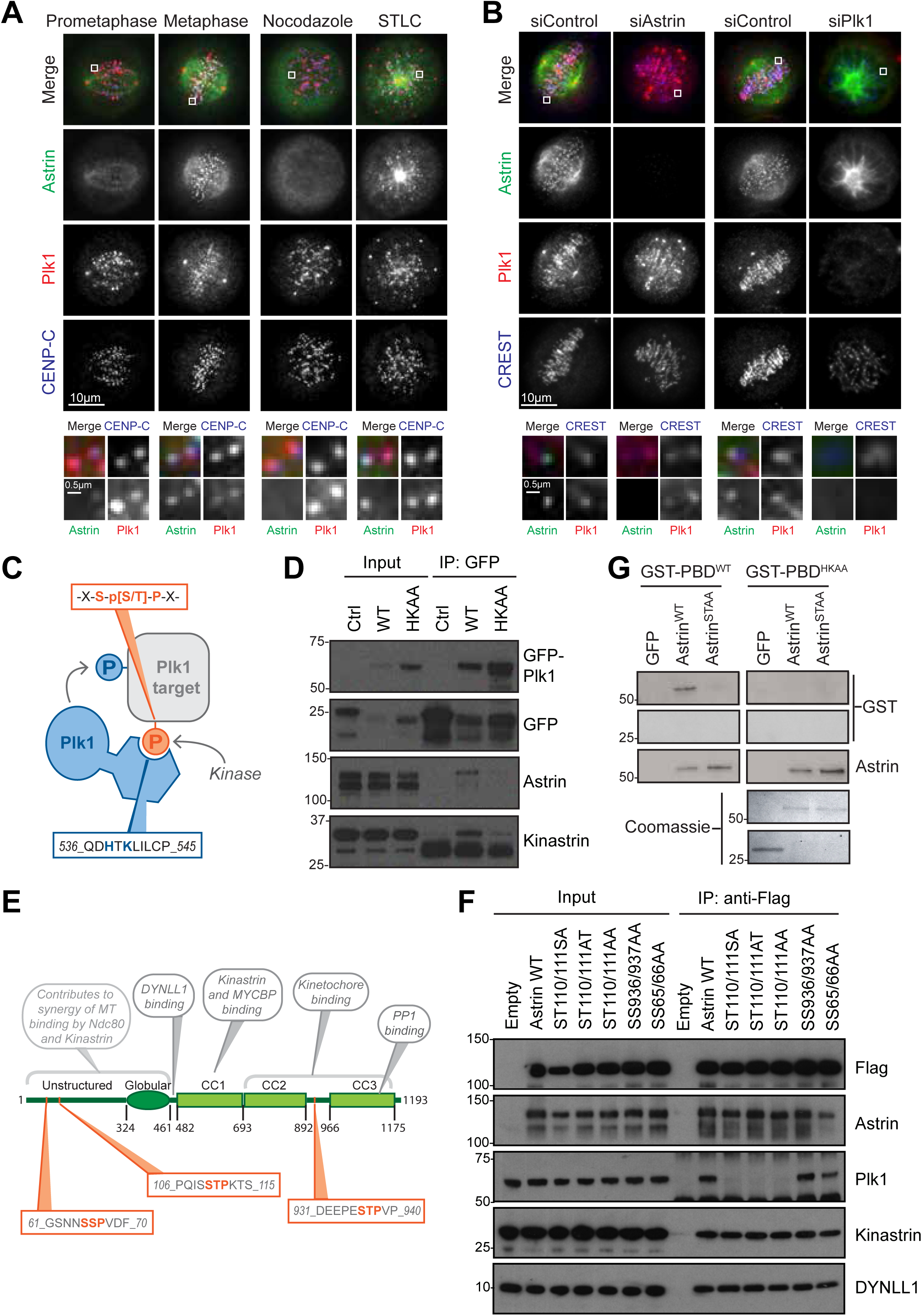
Plk1 binds to a site in the N-terminal domain of Astrin. **A)** Immunofluorescence of Plk1 and Astrin in HeLa cells. Prometaphase and metaphase cells were imaged from an asynchronous cell population; nocodazole and STLC were used to arrest cells in mitosis with unattached and attached kinetochores, respectively. Images were scaled individually to show localisation of lower-intensity Plk1 at attached kinetochores. **B)** Immunofluorescence of Plk1 and Astrin in HeLa cells depleted of Astrin, Plk1 or control (siGL2). **C)** Schematic of Plk1 binding to target motifs via H538 and K540 in the Polo-box domain (PBD). **D)** Plk1 WT and Polo-box domain mutants were transiently expressed and immunoprecipitated from Hek293T cells and analysed by Western blotting. **E)** Schematic of Astrin showing the location of three potential Plk1 binding sites. **F)** Flag-Astrin WT and S-[S/T]-P mutants were immunoprecipitated from Hek293T cells and analysed by Western blotting. **G)** GFP control or Astrin N-terminal domains (aa1-481) were incubated with recombinant Cdk1-Cyclin B1 before Far Western blotting with Plk1 Polo-box domain, WT (left) and binding-deficient mutant HKAA (right).

To confirm the physical association between Astrin and Plk1, reciprocal immunoprecipitations of Astrin and Plk1, respectively, from STLC-treated HeLa cell were carried out. As expected, in Astrin immunoprecipitations Kinastrin was strongly enriched and Plk1 was co-precipitated (Figure S1A). In Plk1 immunoprecipitations, both Astrin and Kinastrin were co-precipitated (Figure S1A). Interestingly, Plk1 only precipitated the slower migrating, mitotically phosphorylated form of Astrin (Chung et al., 2016), suggesting that the interaction between Plk1 and the Astrin complex is phosphorylation-dependent. Plk1 is known to bind to interaction partners via its C-terminal polo-box domain (PBD) to phosphorylated docking sites containing the motif S-pS/pT-P/X (Figure 1C) (Elia et al., 2003a; Elia et al., 2003b). GFP pulldowns from cells transiently expressing GFP-Plk1^WT^ or GFP-Plk1^HKAA^, the latter a version of Plk1 with two key residues required for phospho-specific binding mutated to alanine (Elia et al., 2003b; Hanisch et al., 2006), confirmed that the interaction between Plk1 and Astrin depended on an intact PBD, and that only phosphorylated Astrin interacted with Plk1 (Figure 1D). Astrin contains three potential SSP or STP PBD docking sites at positions 65-67, 110-112 and 936-938 (Figure 1E). In order to identify which of these three motifs is required for Plk1 binding to Astrin, the consecutive serine and threonine residues in all of the motifs were mutated to alanines, and the WT and mutated Astrin versions were then transiently expressed in HEK 293T cells. Pull down of these proteins followed by blotting for Plk1 revealed that mutation of serine and threonine at amino acids 110 and 111 or either one alone resulted in loss of associated Plk1, whereas the equivalent double mutation at residues 936 and 937 and 65 and 66, respectively, had no or only a mild effect on Plk1 binding (Figure 1F). This analysis indicated that the docking site for Plk1 on Astrin is composed of the sequence STP at positions 110 to 112. This site is a canonical CDK1 site and has been reported to be phosphorylated *in vivo* by several mass spectrometry studies (Daub et al., 2008; Hegemann et al., 2011; Kettenbach et al., 2011; Nousiainen et al., 2006). To recapitulate PBD binding to Astrin *in vitro*, recombinant N-terminal fragments of Astrin (amino acids 1-481) containing either the WT amino acid sequence or the ST110/111AA mutation (from hereon referred to as Astrin^STAA^) were phosphorylated in vitro with recombinant CDK1-cyclin B1, immobilised on nitrocellulose membrane by Western blotting and incubated with recombinant GST-tagged PBD. Binding of GST-PBD to membrane-bound N-Astrin was visualised with an anti-GST antibody (Figure 1G). This experiment demonstrated that a successful interaction between CDK1-phosphorylated N-terminal Astrin is dependent on an intact, CDK1-phosphorylated PBD binding motif in Astrin as well as the presence of the critical phospho-site binding residues, His538 and Lys540, in the Plk1-PBD (Elia et al., 2003b) (Figure 1C and 1G).

Taken together, our data indicate that Plk1 associates with Astrin via a single, CDK1-phosphorylated docking motif at amino acid positions 110-112, consistent with the observation that the interaction between Astrin and Plk1 is dependent on CDK1 activity (Kettenbach et al., 2018).

### Plk1 phosphorylates the N-terminus of Astrin

The general idea of Plk1 function through the cell cycle is that Plk1 localises to distinct binding partners via phosphorylated docking motifs which then puts it into the position to phosphorylate the docking partner itself or substrates in the vicinity of the docking partner (Barr et al., 2004; Lera et al., 2016). This mode of operation confers exquisite spatial and temporal control to substrate selection by Plk1. The docking of Plk1 to Astrin suggested that Astrin, or another member of the Astrin complex, may be a Plk1 target. This idea was corroborated by the analysis of the electrophoretic running properties of Astrin in lysates obtained from cells arrested in mitosis by different treatments (Figure 2A). Confirming previous reports, Astrin was strongly upshifted, indicative of phosphorylation, in nocodazole arrested mitotic cells when compared to thymidine-arrested, interphase cells (Chung et al., 2016). This upshift was further enhanced by arresting cells in STLC, mimicking conditions under which Plk1 and Astrin co-localise at the kinetochore (Figure 1A). Interestingly, this additional upshift was reversed to the nocodazole level by incubating STLC-arrested cells with the Plk1 inhibitor BI2536 for 30 mins prior to cell lysis (Lenart et al., 2007) (Figure 2A, compare lanes 2, 3 and 4), suggesting that the additional upshift observed in STLC arrested cells was due to Plk1 phosphorylation. To investigate this possibility in more detail, Astrin was immunoprecipitated from HeLa cells that had been mitotically arrested with STLC and mock-treated or treated with BI2536 for 30 mins prior to cell harvest. The phosphorylation status of the Astrin-Kinastrin complex was then analysed by mass spectrometry. This analysis revealed that, at least under these conditions, only Astrin, but not any other part of the Astrin-Kinastrin complex, was phosphorylated by Plk1 on four sites in the N-terminus of the protein (S157, S159, S353 and S411) (Figure 2B and 2C, Figure S2A). These sites were prominent phospho-sites in the mock-treated Astrin immunoprecipitations but reduced in intensity after BI2536 treatment by at least 2-fold (Figure 2B, Figure S2A) while the docking motif phospho-Thr111 was detected but the intensity not changed by PLk1 inhibition (Figure 2B). All four sites are conserved in most mammalian species (Figure 2C). Of the identified sites S353 fits well with the Plk1 consensus phosphorylation motif (E/N/D(Q))X(S/T)Φ identified in other studies (Figure 2C) (Dou et al., 2011; Kettenbach et al., 2011; Nakajima et al., 2003; Santamaria et al., 2011) which indicated that a polar or acidic residue in position -2 promotes Plk1 phosphorylation. S157/S159 both have a phosphorylatable S/T in position -2, giving rise to the interesting possibility that by phosphorylating S157 Plk1 may create its own consensus site at S159. S411 could not be assigned with absolute certainty as the phosphorylated residue because of the presence of two S/T residues following S411 (Figure S2A).

**Figure 2.**
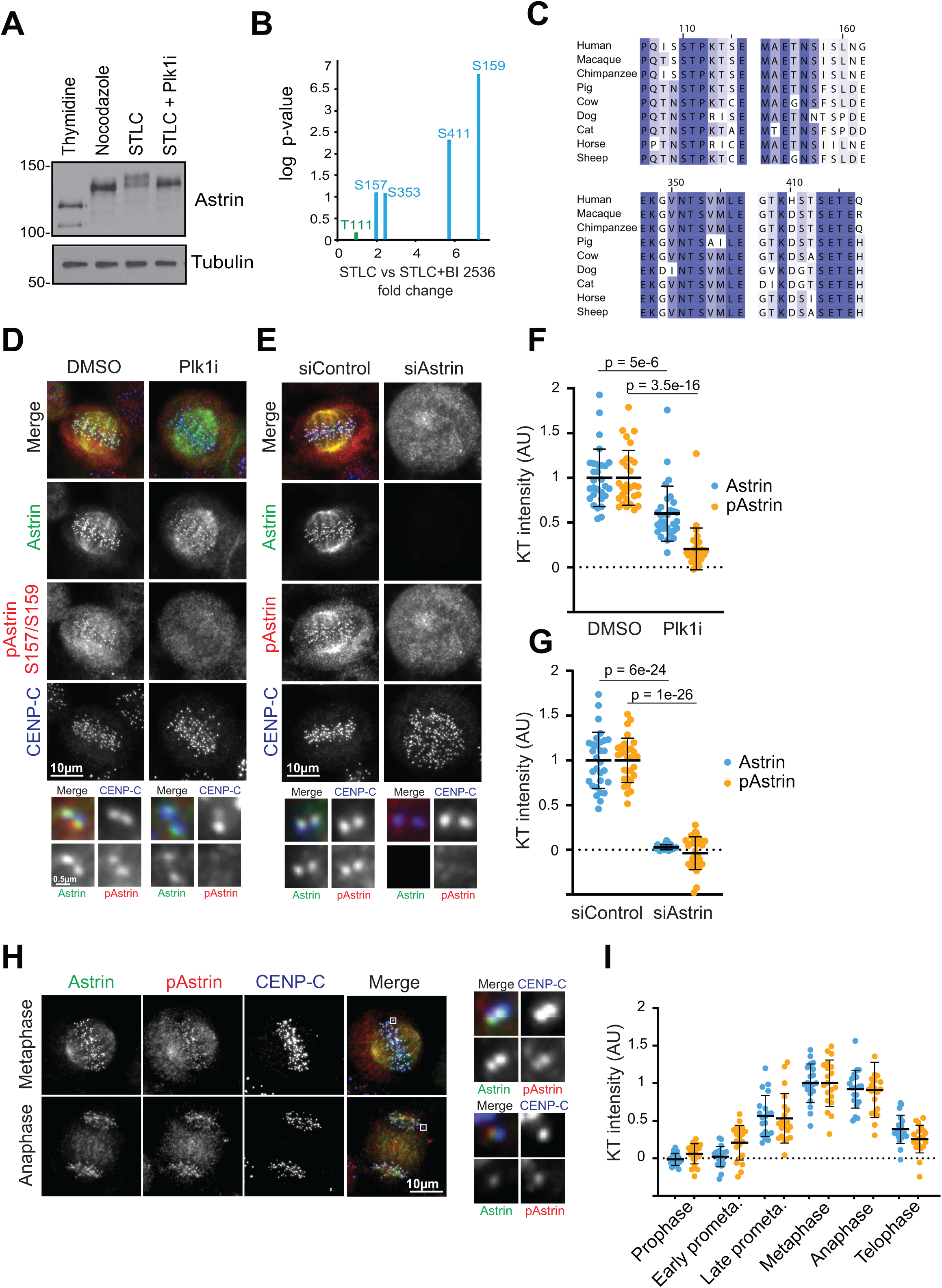
Plk1 phosphorylates the N-terminal domain of Astrin in mitosis. **A)** Cells arrested in different cell cycle states were lysed and Astrin electrophoretic mobility analysed by Western blotting. **B)** Astrin was immunoprecipitated from HeLa cells arrested in STLC with and without a 30 min treatment with Plk1 inhibitor BI 2536, and the phospho-proteome analysed by mass spectrometry. **C)** Clustal alignments of peptide sequences surrounding the Plk1 binding and target sites in the Astrin N-terminal domain. **D)** A phospho-specific antibody was raised against the dually phosphorylated Astrin peptide pS157-pS159. Immunofluorescence imaging of pAstrin pS157-pS159 in HeLa cells following 30 min treatment with Plk1 inhibitor or DMSO control. **E)** Immunofluorescence imaging of pAstrin pS157-pS159 in control- and Astrin-depleted HeLa cells. **F)** and **G)** Quantitation of kinetochore intensities for D) and E) respectively. Each dot represents the average intensity of 20 kinetochores from a single cell. 10 cells were quantified per condition in 3 independent repeats. Lines represent Mean ±SD. **H)** Immunofluorescence imaging of pAstrin pS157-pS159 in asynchronous HeLa cells. **I)** Quantitation the kinetochore intensity of Astrin and pAstrin measured relative to CENP-C kinetochore intensity, and normalised to maximum intensity (at metaphase; =1). Each dot represents the average intensity of 20 kinetochores from a single cell. 10 cells were quantified per mitotic stage in 2 independent repeats. Lines represent Mean ±SD. All p-values were calculated by two-tailed Student t test.

Antibody staining with a highly specific antibody raised against the doubly phosphorylated pS157/pS159 peptide (pAstrin), showed that these antibodies prominently stained kinetochores, and that the staining was dependent on the presence of both Astrin and Plk1 activity (Figure 2D-2G). An antibody raised against pS353 gave a qualitatively similar kinetochore staining (Figure S2B). However, this antibody also displayed spindle background staining which was Plk1 but not Astrin dependent, and this antibody was therefore not used for further analysis.

Our data suggest that the co-localisation of Astrin and Plk1 at kinetochores may lead to the interaction of the two proteins and result in Astrin phosphorylation by Plk1. When we followed pAstrin staining through the cell cycle, the phospho-specific signal largely followed the total Astrin staining (Figure 2H and 2I). Interestingly, pAstrin staining, like total Astrin, was retained on anaphase kinetochores, suggesting that these phospho-sites were not turned over upon anaphase onset.

### Phosphorylation of Astrin, but not spindle bipolarity, is dependent on the Plk1-Astrin association

Depletion of Astrin results in a range of mitotic abnormalities including the formation of multipolar spindles, impaired chromosome congression and spindle checkpoint mediated mitotic arrest (Dunsch et al., 2011; Gruber et al., 2002; Thein et al., 2007). To test whether the association between Plk1 and Astrin was relevant for the functionality of the Astrin-Kinastrin complex, RNAi rescue assays using HeLa-Flp-in cells inducibly expressing GFP-Astrin^WT^ or the GFP-Astrin^STAA^ Plk1-binding mutant were employed (Figure 3A). In these cells, expression levels of transgenic Astrin were comparable between cell lines and to the endogenous protein (Figure 3B). Depletion of Astrin by RNAi results in a mitotic arrest with large numbers of multipolar cells (Dunsch et al., 2011; Gruber et al., 2002; Thein et al., 2007). Replacement of endogenous Astrin with GFP-Astrin led to a rescue of the Astrin phenotype indicated by normal progression through the cell cycle as judged by live cell imaging (Figure S3D and S3E) and by the significantly reduced number of multipolar spindles in comparison to the non-rescued RNAi situation (Figure 3C and 3E). Interestingly, RNAi rescue with GFP-Astrin^STAA^ restored Astrin functionality to a similar extent as GFP-Astrin^WT^ indicating that the association of Plk1 with the Astrin-Kinastrin complex was not necessary for the Astrin-Kinastrin mediated formation and maintenance of a bipolar spindle, or cell cycle progression (Figure 3C and 3E, Figure S3D and S3E). These data are in line with the idea that the entire N-terminus of Astrin may be dispensable for Astrin’s function in promoting spindle bipolarity (Kern et al., 2017). Indeed, our own analysis confirmed that expression of GFP-Astrin lacking the first 464 amino acids (GFP-Astrin^ΔN^) in HeLa cells depleted of endogenous Astrin rescued spindle bipolarity and progress through an unperturbed cell cycle (Figures S3B-E).

**Figure 3.**
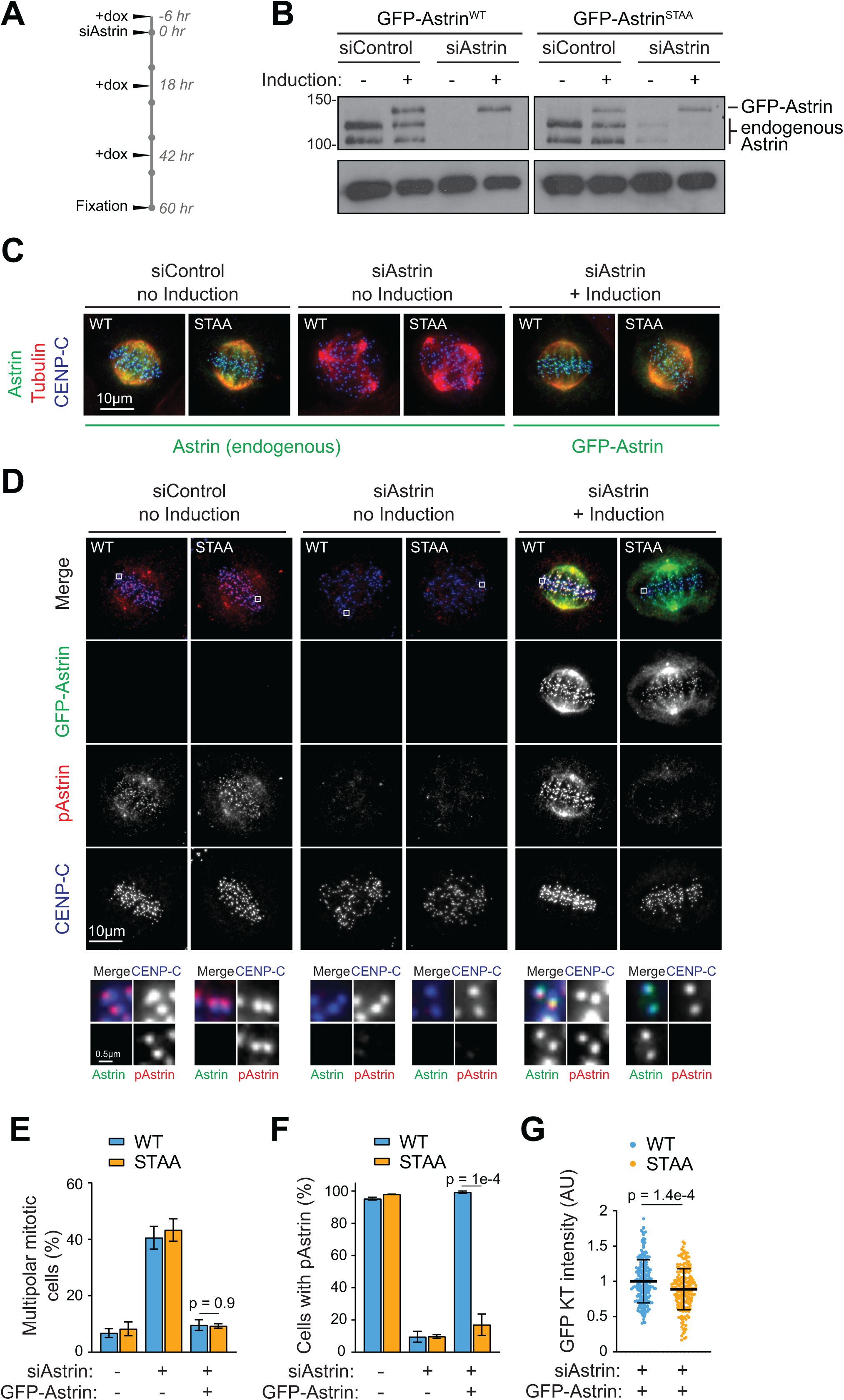
The Plk1 binding site is required for Astrin phosphorylation but not for spindle bipolarity. **A)** Schematic of timeline for Astrin RNAi rescue experiment. **B)** HeLa Flp-In TRex cells were depleted of endogenous Astrin and induced to express GFP-Astrin^WT^ or GFP-Astrin^STAA^ as shown in A). Astrin was analysed by immunoblotting. **C)** Immunofluorescent images of HeLa Flp-In TRex cells depleted of endogenous Astrin and induced to express GFP-Astrin^WT^ or GFP-Astrin^STAA^. **D)** The phosphorylation status of Astrin S157/9 sites was analysed by immunofluorescence in cells treated as in B) and C). **E)** The percentage of multipolar mitotic cells was counted from the conditions shown in C). Bars represent the Mean ±SEM of 4 independent experiments, with 50-150 cells counted per condition per repeat. **F)** Quantitation of the experiment shown in D). Cells with clearly visible GFP-Astrin at kinetochores were assessed as positive or negative for pAstrin kinetochore staining. Bars represent the Mean ±SEM of 3 independent experiments, with 30-100 cells counted per condition per repeat. **G)** Quantitation of GFP-Astrin kinetochore intensities. Cells treated as in A) were fixed in PFA to preserve any cytoplasmic pool of GFP-Astrin and the intensities of the whole cell measured to ensure equivalent expression. Each dot represents an individual kinetochore; bars indicate Mean ±SD. All p-values were calculated by two-tailed Student t test.

However, when we analysed Plk1-induced phospho-Astrin staining in mitotic cells depleted of endogenous Astrin and rescued with either GFP-Astrin^WT^ or GFP-Astrin^STAA^, we found that Plk1 phosphorylation of Astrin was strictly dependent on the presence of the Plk1-binding site in Astrin and was not restored in Astrin depleted cells expressing GFP-Astrin^STAA^ (Figure 3D-3F). This suggested that direct docking of Plk1 to Astrin was required for the phosphorylation of Astrin by Plk1, and that this could not be carried out by Plk1 bound to other interactions partners at the kinetochore, although, as expected, Plk1 was still present at kinetochores when binding to Astrin was prevented (Figure S3A). Furthermore, we noticed that Astrin kinetochore localisation was slightly impaired in cells expressing GFP-Astrin^STAA^, suggesting that the interaction of Astrin with Plk1 promotes Astrin kinetochore localisation (Figure 3G).

### Phosphorylation of the Astrin N-terminal domain by Plk1 contributes to kinetochore-microtubule attachment stability

Taken together, our data suggested that there may be a function of Astrin-associated Plk1 that was not captured by assessing spindle bipolarity or cell cycle progression under unperturbed conditions. We hypothesised that a possible function of Plk1 in promoting localisation of the Astrin-Kinastrin complex to kinetochores may only become necessary under conditions where microtubule-kinetochore are put under additional stress, e.g. in situations of prolonged mitotic arrest. In order to address this possibility, we turned to situations where normal cell cycle progression was impeded. First, we analysed the ability of different Astrin mutants to localise to kinetochores in cells arrested with a monopolar spindle induced by STLC treatment (Figure 4A). Under these conditions, kinetochores are syntelically oriented and lack tension, and for most kinetochore pairs, only one kinetochore is attached to microtubules. In these spindles, GFP-Astrin^WT^ was localised on average to 60.5 ± 10.8% of kinetochores whereas GFP-Astrin^STAA^ only localised to 46.8 ± 12.4% of kinetochores (Figure 4B). Strikingly, GFP-Astrin^ΔN^ lacking the entire N-terminus was only found at 27.7 ± 9.1% of kinetochores and even when it was found at kinetochores, displayed a much-reduced intensity (Figure 4B and 4C). To test whether the different versions of Astrin affected the stability of the microtubule-kinetochore attachments in STLC-treated cells, the immunofluorescence analysis of GFP-Astrin was combined with a 9 min cold treatment (Figure 4A-C). Under these conditions, only stable microtubule-kinetochore attachments are preserved (Rieder, 1981). In this situation, the contribution of the Astrin N-terminus to kinetochore localisation and stability of attachments was particularly evident, as in cold-treated STLC spindles GFP-Astrin^ΔN^ was completely lost from kinetochores (Figure 4A, right panel) while GFP-Astrin^WT^ and, with lower efficiency, GFP-Astrin^STAA^, still localised to kinetochores. Furthermore, versions of Astrin in which the four Plk1 sites (S157, S159, S353, S411) as well as the two phosphorylatable serine/threonines following S411 (T412, S413) had been mutated to alanine (GFP-Astrin^6A^), showed a similarly impaired kinetochore localisation as GFP-Astrin^STAA^, whereas the phospho-mimetic mutant GFP-Astrin^6D^ behaved like wild type (Figure S4B and S4C). Taken together, these data suggest that the presence of the Astrin N-terminus, and within the N-terminus the association with Plk1, promotes Astrin localisation to kinetochores.

**Figure 4.**
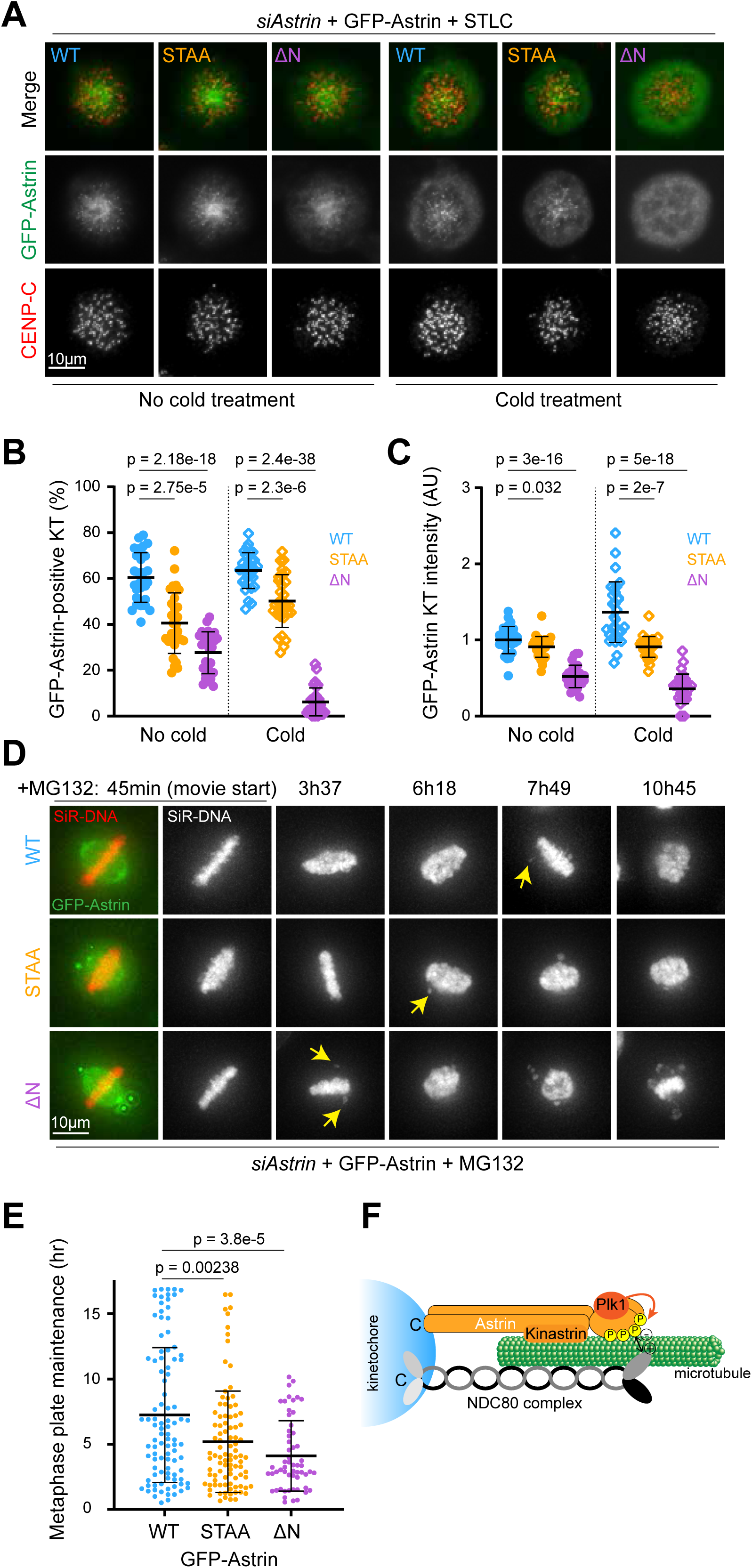
Phosphorylation of the Astrin N-terminal domain by Plk1 contributes to kinetochore-microtubule attachment stability. **A)** HeLa Flp-In TRex cells depleted of endogenous Astrin and induced to express GFP-Astrin^WT^, GFP-Astrin^STAA^, or GFP-Astrin^ΔN^ were arrested overnight with STLC. Cells were either fixed directly or subjected to 9 minutes cold treatment immediately prior to fixation.**B)** Quantitation of cells as in A). The number of Astrin-positive kinetochores was counted and calculated as a percentage of visible kinetochores from CENP-C staining. For each condition 30 cells were analysed from 2 or 3 independent repeats. Each dot represents a single cell; error bars show Mean ±SD. **C)** The intensity of GFP-Astrin-positive kinetochores was measured for the cells analysed in B). 20 Astrin-positive kinetochores were measured per cell (apart from the GFP-Astrin^ΔN^ cells which had fewer than 20 Astrin-positive kinetochores). Each dot represents the mean kinetochore intensity of an individual cell; error bars show Mean+-SD. **D)** HeLa Flp-In TRex cells were depleted of endogenous Astrin and induced to express GFP-Astrin^WT^, GFP-Astrin^STAA^, or GFP-Astrin^ΔN^. The ability of cells to maintain a metaphase plate in MG132 arrest was analysed by live cell imaging. Yellow arrows indicate the first incidence of chromosome loss from the metaphase plate. **E)** The time from MG132 addition to first visible loss of chromosomes from the metaphase plate was measured for cells as in D). Each dot represents an individual cell. Cells are from 3 or more independent repeats; error bars show Mean ±SD. All p-values were calculated by two-tailed Student t test. **F)**. Model illustrating how Plk1-phosphorylation of the Astrin N-terminal domain promotes binding of Astrin to NDC80 to stabilize microtubule-kinetochore attachments. For simplicity, Astrin binding partners DYNNL1 and MYCBP are not depicted.

These observations prompted us to investigate whether the Astrin-Plk1 complex was required for the maintenance of stable microtubule-kinetochore attachments under conditions of prolonged mitotic arrest. To test this, mitotic cells expressing the different GFP-Astrin variants were treated with the proteasome inhibitor MG132 to prevent anaphase onset, and assessed for their ability to stably maintain a metaphase plate for extended periods of time (Figure 4D and 4E). Cells expressing GFP-Astrin^WT^ upheld a metaphase plate on average for 434 ± 310 min before chromosomes started leaving the plate. The ability to maintain a metaphase plate was reduced to 311± 234 min in the GFP-Astrin^STAA^ mutant and even further to 246 ± 161 min in GFP-Astrin^ΔN^. Together with the localisation data in Figure 4B, these data suggest that the phosphorylated N-terminus of Astrin promotes accumulation of the Astrin complex at attached kinetochores and consequent stabilisation of microtubule-kinetochore attachments. Our data suggest that this becomes important when microtubule-kinetochore attachments have to be maintained for longer periods of time (Figure 4D and 4E).

### Astrin and Plk1 promote microtubule-kinetochore attachments synergistically

The formation of stable microtubule-kinetochore attachments is of critical importance for the faithful execution of chromosome segregation. Both the Astrin-Kinastrin complex and the Plk1 kinase have been implicated in promoting the formation of stable microtubule-kinetochore attachments but whether these two factors synergise has so far not been investigated. Our data here show that Plk1 interacts with Astrin via a canonical PBD binding site in the globular N-terminus of Astrin and deposits phosphorylations on at least four sites in the same part of the molecule. It should be noted that in addition to the PBD binding site and the Plk1 phosphorylation sites that we mapped here, the N-terminus of Astrin also contains another six canonical CDK1 phosphorylation sites. The high incidence of phosphorylation sites in this part of the Astrin molecule explains the striking electrophoretic upshift that is observed for Astrin isolated from mitotic cells (Figure 2A). We found that the full mitotic upshift, requiring both CDK1 and Plk1 phosphorylation, was only achieved in the presence of microtubules (Figure 2A). Although Plk1 is highly enriched on unattached kinetochores, Astrin is not found on kinetochores in the absence of microtubules and thus cannot interact with Plk1 efficiently in this situation (Figure 1A). It has previously been shown that the C-terminus of Astrin contains the main kinetochore binding site and that the N-terminus alone does not localise to any spindle structures (Conti et al., 2019; Dunsch et al., 2011; Kern et al., 2017). Interestingly, though, in biochemical binding assays the Astrin N-terminus promoted the interaction of the Astrin-Kinastrin complex with purified NDC80 complex (Kern et al., 2017). This is consistent with our immunofluorescence-based results which indicated that the presence of the Astrin N-terminus and the PBD-binding motif within the N-terminus facilitated the recruitment of the Astrin complex to kinetochores and thus promoted the formation of cold-stable microtubule-kinetochore attachments (Figure 4B). How the highly negatively charged N-terminal domain in Astrin supports the recruitment of the Astrin-complex to bioriented kinetochores is currently not clear. It has been reported that localisation of the Astrin-complex to kinetochores is negatively regulated by Aurora B phosphorylation (Manning et al., 2010; Schmidt et al., 2010), and an interaction between the positively charged Aurora B substrate recognition motifs in their unphosphorylated forms and the negatively charged Astrin N-terminus may in part explain this behaviour.

Our data suggest that the additional contribution of the Astrin N-terminal domain to kinetochore binding becomes particularly important in situations when mitotic progression is delayed, in our assay mimicked by a mitotic arrest imposed by MG132 treatment. Physiologically, this may be more important in meiotic rather than mitotic cell divisions, since mammalian oocytes have to be able to maintain a prolonged arrest in metaphase of meiosis II (Madgwick and Jones, 2007) which may require additional mechanisms to stabilise microtubule-kinetochore attachments during this period. Further analysis of the requirement of the Astrin-Kinastrin complex in oocyte meiosis may thus be informative.

## Acknowledgements

ZG and PU were supported by a Medical Research Council Senior Non-Clinical Fellowship awarded to UG (MR/K006703/1) and a Biotechnology and Biological Sciences Research Council Strategic LoLa grant (BB/M00354X/1). CB was supported by a Medical Research Council studentship (1512121). We acknowledge Kerstin Thein and Anja Dunsch for initial observations, Sabine Hiltscher for help with recombinant protein production, James Bancroft for assistance with live cell imaging experiments and Shabaz Mohammed for help with mass spectrometry.

## Material and Methods

### Chemicals and antibodies

General laboratory chemicals and reagents were obtained from Sigma-Aldrich and Thermo-Fisher Scientific. Drugs were dissolved in DMSO unless specifically indicated. Inhibitors were obtained from Sigma Aldrich (Eg5 inhibitor STLC, 10 mM stock), Selleck (Plk1 inhibitor BI2536, 2 mM stock), Insight Bioscience (proteasome inhibitor MG132 20 mM stock) and Merck (microtubule polymerisation inhibitor nocodazole 0.66 mM stock). Thymidine (Sigma Aldrich, 100mM stock) and doxycycline (Invivogen, 2 mM stock) were dissolved in water.

Commercially available polyclonal (pAb) or monoclonal (mAb) antibodies were used for Tubulin (Mouse mAb; Sigma, [DM1A] T6199), Plk1 (Mouse mAb; Santa Cruz clone F-8, sc-17783). Antibodies against Astrin and Kinastrin/SKAP have been described before (Dunsch et al., 2011; Thein et al., 2007). Antibodies against the doubly phosphorylated Astrin p157/159 peptide CMAETN(pS)I(pS)LNGP and pS353 peptide CKGVNTSVMLEN, coupled to KLH, were raised in sheep (Orygen Antibodies). The antibodies were affinity-purified from the serum on the immobilised immunising peptide.

Secondary donkey antibodies against mouse, rabbit, guinea pig or sheep and labelled with Alexa Fluor 488, Alexa Fluor 555, Alexa Fluor 647, Cy5, or HRP were purchased from Molecular Probes and Jackson ImmunoResearch Laboratories, Inc., respectively. Affinity purified primary and HRP-coupled secondary antibodies were used at 1 µg/ml final concentration. For western blotting, proteins were separated by SDS-PAGE and transferred to nitrocellulose using a Trans-blot Turbo system (Bio-Rad). Protein concentrations were measured by Bradford assay using Protein Assay Dye Reagent Concentrate (Bio-Rad). All western blots were revealed using ECL (GE Healthcare).

### Molecular biology and siRNA reagents

Astrin mutant expression constructs were made using pcDNA5/FRT/TO vectors (Invitrogen) modified to encode the EGFP or FLAG reading frames. Mutagenesis to introduce phospho-site mutations and resistance to Astrin siRNA was performed using the QuikChange method (Agilent Technologies). DNA primers were obtained from Invitrogen. For the knock down of Plk1 small interfering RNA (siRNA) duplexes AACGAGCTGCTTAATGACGAGTT were used (ThermoFisher). For Astrin siRNA oligo #367 (5′-TCCCGACAAC TCACAGAGAAA -3′)(Qiagen) was used as described (Thein et al., 2007).

### Cell culture and transfection

HeLa cells were cultured in DMEM with 1% [vol/vol] GlutaMAX (Life Technologies) containing 10% [vol/vol] bovine calf serum at 37°C and 5% CO_2_. HeLa Flp-In TRex cells expressing GFP-Astrin variants cells were maintained in medium supplemented with 200μg/ml Hygromycin B (Invivogen) and 4μg/ml Blasticidin (Invivogen). For plasmid transfection and siRNA transfection, Mirus LT1 (Mirus Bio LLC) and Oligofectamine (Invitrogen), respectively, were used. HeLa cell lines with single integrated copies of the desired transgene were created using the T-Rex doxycycline-inducible Flp-In system (Invitrogen)(Tighe et al., 2004).

### Immunofluorescence analysis

Cells were fixed with PTEMF fixation buffer (20mM Pipes-KOH, pH 6.8, 0.2% Triton X-100, 1mM MgCl2, 10mM EGTA and 4% formaldehyde) for 12mins at room temperature (RT) (Dunsch et al., 2011). Coverslips were washed 3x with PBS before transfer into blocking solution (PBS with 3% BSA) and blocking proceeded for 30mins. Primary and secondary antibody incubations were performed in blocking solution for 1hr and 30mins respectively. DNA was stained in the secondary antibody incubation with 1μg/ml DAPI. Coverslips were washed 3x with PBS after each incubation. Coverslips were dried and mounted in moviol on microscope slides.

### RNAi rescue experiments

For Astrin siRNA rescue experiments, HeLa Flp-In TRex cells inducibly expressing GFP-Astrin^WT^ or mutants thereof were used. Astrin siRNA rescue was performed by induction with 2 µM doxycycline of GFP-Astrin transgenes resistant to Astrin RNAi, for 6 hours prior to a 60 hr siRNA depletion of endogenous Astrin using oligo #367. A second induction was performed 18 hrs into the siRNA depletion. For live cell imaging, cells were treated with 2 mM thymidine 18 hrs after RNAi addition, for 18 hours. The thymidine was removed by washing 3 times with DMEM, with 2 µM doxycycline re-added in the final wash. SiR-DNA (Spirochrome) was added to the final wash of the thymidine release at a concentration of 100nM, and imaging commenced 9 hours later. Live cell imaging was performed on a Deltavision Elite system using an inverted microscope (IX81; Olympus) and equipped with a QuantEM EMCCD camera (Photometrics). Cell were placed in a 37°C and 5% CO2 environmental chamber (Tokai Hit) on the microscope stage with lens heating collar. Imaging was performed using a 60× NA 1.4 oil immersion objective lens. Cells were imaged using 5% 488 nm laser power with 50 ms exposure for GFP-Astrin and 2% 647 nm laser power with 10 ms exposure for SiR-DNA. 7 axial planes were captured at 2 μm apart (Z then wavelength) at an interval of 2 min for 12h to analyse mitotic progression. For Mg132-arrested live assays, 9 hours after thymidine release MG132 was added to a final concentration of 20 μM immediately before cells were placed on the microscope, and image capture was begun as soon as possible. 11 axial planes were captured at 2 μm apart (Z then wavelength) at an interval of 3 min for 16h. These images were then used to determine when the first chromosomes left the mitotic metaphase plate. For the majority of analysed cells which were already at metaphase when imaging commenced, the set up time for that experiment (from Mg132 addition to start of imaging) was added during analysis; for a minority of cells which entered mitosis during imaging, the time from last chromosome congression to first chromosome loss was measured.

### Far Western blot analysis

1 µg of purified His-tagged N-terminal Astrin (aa1-481) containing WT or ST110/111AA mutations, respectively, was incubated with 10μl 2x MEB (100mM Tris-HCl pH7.4, 100mM KCl, 20mM MgCl2, 40mM Sodium beta-glycerolphosphate, 30mM EGTA), 1μl of 10x ATP/DTT mix (10mM ATP, 100mM DTT), 100ng Cdk1-cyc B1 (Abcam) and made up to 20μl in water. Reactions were incubated for 60mins at 30°C. Samples were denatured in Laemmli buffer, run on 10% SDS-PAGE gels and transferred onto nitrocellulose membrane. The membrane was incubated overnight in blocking (50mM Tris-HCl pH7.5, 137mM NaCl, 0.1% Tween, 4% milk powder). Samples were incubated with 1μg/ ml His-GST-tagged Plk1-PBD constructs (either WT or HKAA mutants) for 8hrs at RT. The membrane was then washed 3x in blocking buffer and incubated overnight with rabbit anti-GST antibody before 3x washes (PBS+0.1% Tween) before incubation with donkey-anti-rabbit secondary antibody coupled to HRP.

### FLAG pull-down assays

Flag-Astrin constructs were transiently transfected into HEK293T cells using TransIT-LT1 transfection reagent (Mirus Bio) according to the manufacturer’s protocol. 24 hrs post-transfection cells were arrested with 10 µM STLC for 14hrs overnight. Cells were collected via a mitotic shake-off and lysed in 50 mM Tris-HCl, pH8.0, 150 mM NCL, 1% IGEPAL, protease inhibitor cocktail (Sigma) and phosphatase inhibitor cocktail 3 (Sigma) prior to immunoprecipitation with FLAG agarose beads (Sigma).

### Immunoprecipitations

For the analysis of Astrin phosphorylation sites, HeLa cells stably expressing GFP-Astrin (Dunsch et al., 2011) were arrested overnight in STLC. One half of the cells was treated for 30 mins with Plk1 inhibitor BI2536 prior to harvest by mitotic shake-off; the other half was left untreated. Cell pellets were lysed (50mM HEPES pH8, 100mM KCl, 1%IGEPAL, 0.25% Triton X-100, 1mM DTT, 1:250 Protease Inhibitor Cocktail (Sigma), 1:100 Phosphatase Inhibitor cocktail 3 (Sigma), 50mM EDTA, 1mM PMSF, 50U micrococcal nuclease) for 30mins at 4°C before centrifugation (14,000rpm for 15 min). Per 1mg of cell lysate 1μg of the appropriate antibody Sheep anti-GFP antibody was added. Immunoprecipitated proteins were eluted first with 0.1 M glycine, pH 2.6 followed by 50 mM Tris-HCl, Ph 8.5, 8M urea. Western blots were used to confirm the success of the immunoprecipitations.

### Mass spectrometry

For mass spectrometry analysis samples were processed using filter aided sample preparation (FASP) columns. Samples were reduced using 10mM Tris(2-carboxyethyl)phosphine hydrochloride (TCEP) for 30 min followed by alkylation, using 20mM chloroacetaldehyde (CAA), in the dark for 30 min. Proteins were digested with 1.5μg Trypsin (Promega) at 37°C for 12 hours. Samples were reduced to approx. 50μl using a Thermo Scientific SpeedVac concentrator centrifuge. Phospho-peptide enrichment was performed using titanium dioxide microspin columns (TopTip; Glygen). All spin steps were performed at 550 rpm, equivalent to 34 *g*, for 5 min at room temperature. Columns were stripped with 50μl elution buffer (5% ammonia solution in water) and washed 3x with 65μl loading buffer (1M glycolic acid, 80% acetonitrile (ACN), 5% trifluoroacetic acid (TFA)). Samples were diluted 1:1 using concentrated loading buffer (10% TFA, 2M Glycolic acid, 80% ACN) and loaded onto the column 65μl at a time. Following loading, columns were washed once each with loading buffer, wash buffer (0.2% TFA acid in 80% ACN) and finally 20% ACN. Phospho-peptides were eluted using 2x 10 μl of elution buffer into an Eppendorf containing 20μl 5% Formic Acid. Liquid chromatography was performed using an EASY-nano-LC 1000 system (Proxeon). Peptides were loaded onto a 75μm internal diameter guard column (packed with Reprosil-Gold 120 C’8, 3 μm, 120 Å pores) using Solvent A (0.1% [vol/vol] formic acid in water) at 500 bar. Peptide separation was performed by an EASY-Spray column at 45 °C (50 cm × 75 µm ID, PepMap RSLC C18, 2 µm; Thermo Fisher Scientific). A 30 min linear 8-30% [vol/vol] ACN gradient was used with a flow rate of 200 nl/min. An EASY-Spray nano-electrospray ion source was used to introduce peptides into a Q-Exactive mass spectrometer and spectra were acquired with an m/z range of 350-1500. The 20 most abundant peaks were fragmented using CID. MaxQuant (Tyanova et al., 2016a) was used to process the data, and Perseus was used to analyse the mass spectrometry data sets (Tyanova et al., 2016b).

