## Supplemental material for "The association of Plk1 with the Astrin-Kinastrin complex promotes formation and maintenance of a metaphase plate"

#### Supplemental figures

#### Supplemental Figure legends

**Figure S1. Astrin and Plk1 interact biochemically. A)** Immunoprecipitations of Astrin (left) and Plk1 (right) from STLC-arrested HeLa cells were analysed by Western blotting with the indicated antibodies.

**Figure S2. Plk1 phosphorylates four sites in the N-terminus of Astrin. A)** Four Plk1 phosphorylation sites in Astrin were identified by mass spectrometry. **B)** A phospho-specific antibody was raised against Astrin pS353. pAstrin pS353 was analysed by immunofluorescence imaging in HeLa cells. Left panel shows staining following 30min treatment with Plk1 inhibitor or DMSO control; right panel shows staining in control- and Astrin-depleted cells.

**Figure S3. The N-terminus of Astrin is not required for promoting spindle bipolarity. A)** In HeLa Flp-In TRex cells depleted of endogenous Astrin and induced to express GFP-Astrin$^{WT}$ or GFP-Astrin$^{STAA}$, Plk1 kinetochore localization was visualised by immunofluorescence analysis. **B)** HeLa Flp-In TRex cells were depleted of endogenous Astrin and induced to express GFP-Astrin$^{WT}$ or GFP-Astrin$^{\Delta N}$, and stained for tubulin. **C)** The percentage of multipolar mitotic cells was counted for the conditions shown in B). Bars represent the Mean ±SEM of 3 independent experiments, with 50-150 cells counted per condition per repeat. P-value was calculated by two-tailed Student t test. **D)** HeLa Flp-In TRex cells depleted of endogenous Astrin and induced to express GFP-Astrin$^{WT}$, GFP-Astrin$^{STAA}$ or GFP-Astrin$^{\Delta N}$ were analysed by live cell imaging. Images were captured every 2 minutes and the time from nuclear envelope breakdown (NEBD) to anaphase onset was calculated. Each dot represents an individual cell; cells are from 4 (WT) or 2 (STAA, ΔN) independent experiments. **E)** Representative images of cells analysed in D).

**Figure S4. A) Phospho-mimetic mutations in the N-terminus of Astrin promote the kinetochore localisation of Astrin.** HeLa Flp-In TRex cells depleted of endogenous Astrin and induced to express GFP-Astrin$^{WT}$, GFP-Astrin$^{6A}$, or GFP-Astrin$^{6D}$ were arrested overnight with STLC. Cells were either fixed directly or subjected to 9 min cold treatment immediately prior to fixation. **B)** Quantitation of cells as in A). The number of Astrin-positive kinetochores was counted and calculated as a percentage of visible kinetochores from CENP-C staining. For each condition 30 cells were analysed from 2 or 3 independent repeats. Each dot represents a single cell; error bars show Mean ±SD. **C)** The intensity of GFP-Astrin-positive kinetochores was measured for the cells analysed in B). 20 Astrin-positive kinetochores were measured per cell. Each dot represents the mean kinetochore intensity of an individual cell; error bars show Mean ±SD. All p-values were calculated by two-tailed Student t-test.

### Figure S1

A

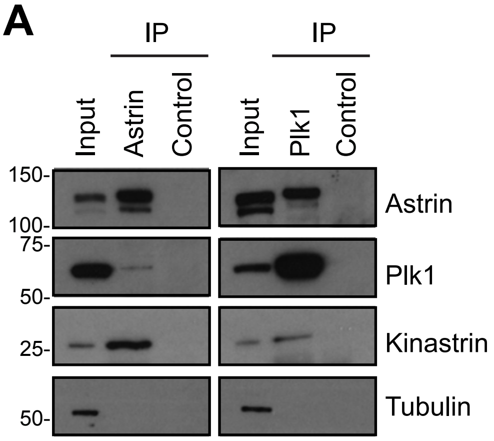
IP
IP
Input
Astrin
Control
Input
Plk1
Control
150-
100-
75-
50-
25-
50-
Astrin
Plk1
Kinastrin
Tubulin

### Figure S2

## A

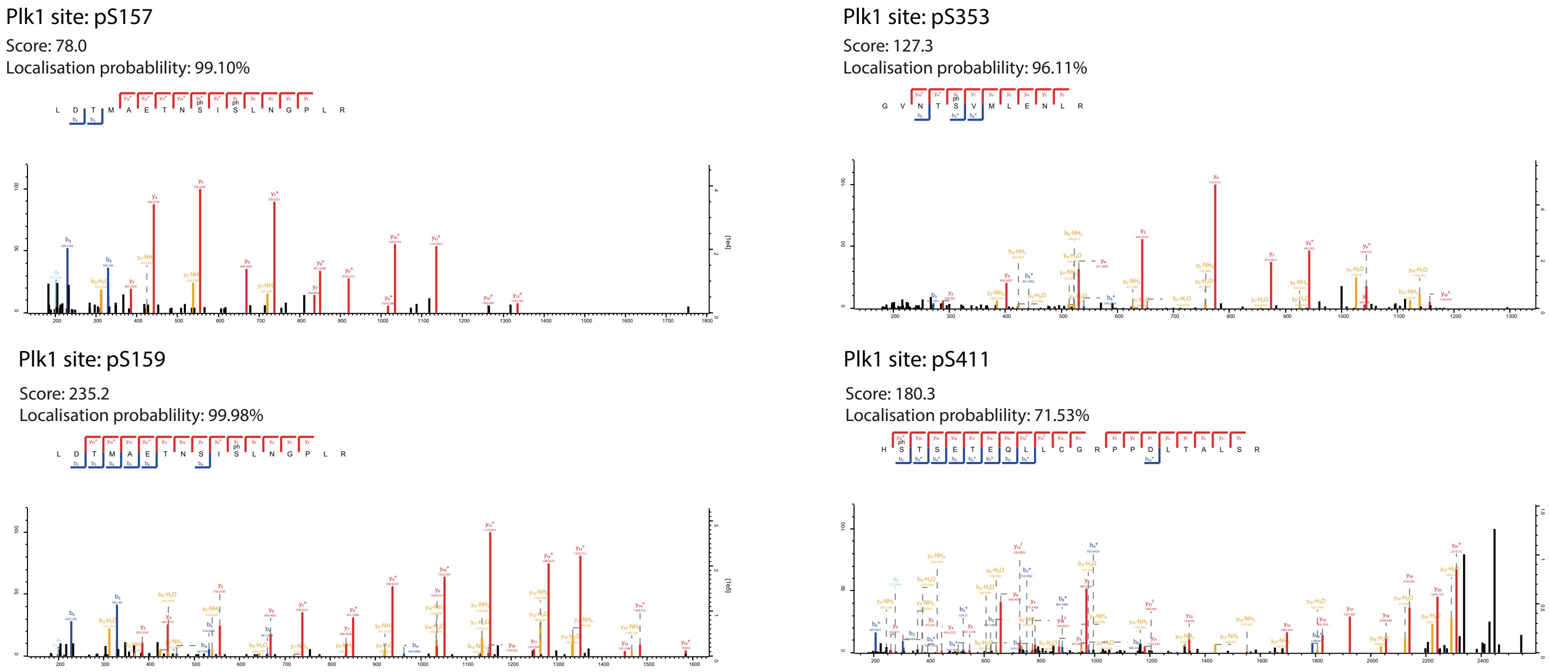

Plk1 site: pS157
Score: 78.0
Localisation probablility: 99.10%
Plk1 site: pS353
Score: 127.3
Localisation probablility: 96.11%
Plk1 site: pS159
Score: 235.2
Localisation probablility: 99.98%
Plk1 site: pS411
Score: 180.3
Localisation probablility: 71.53%


## B

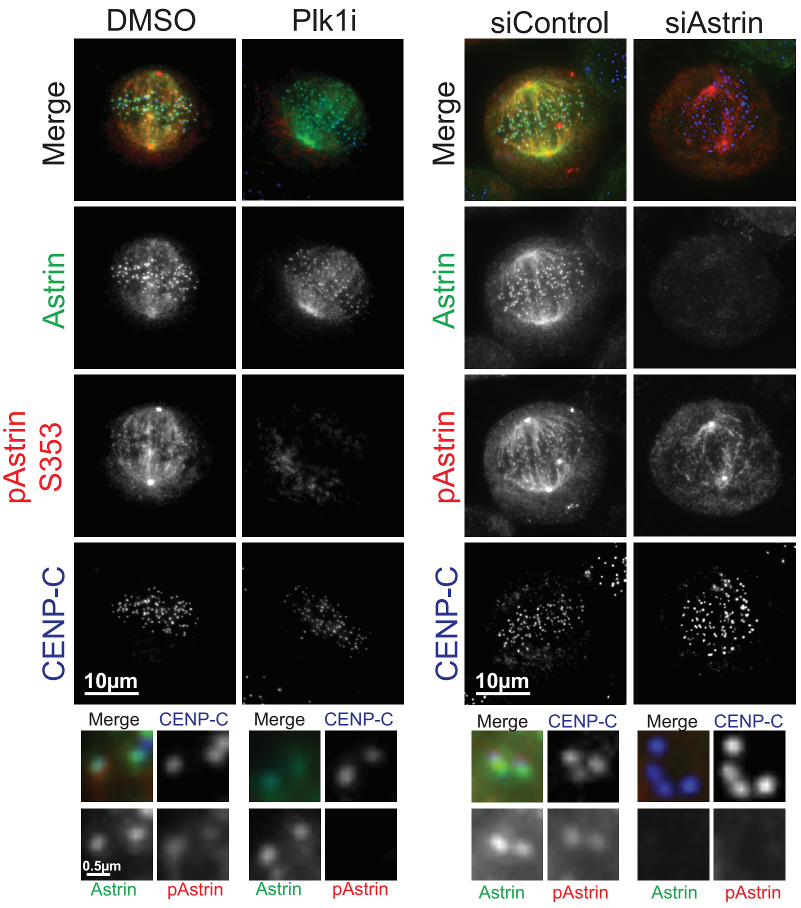

DMSO
Plk1i
siControl
siAstrin
Merge
Astrin
pAstrin S353
pAstrin
CENP-C
10µm
0.5µm
Merge
CENP-C
Astrin
pAstrin

### Figure S3

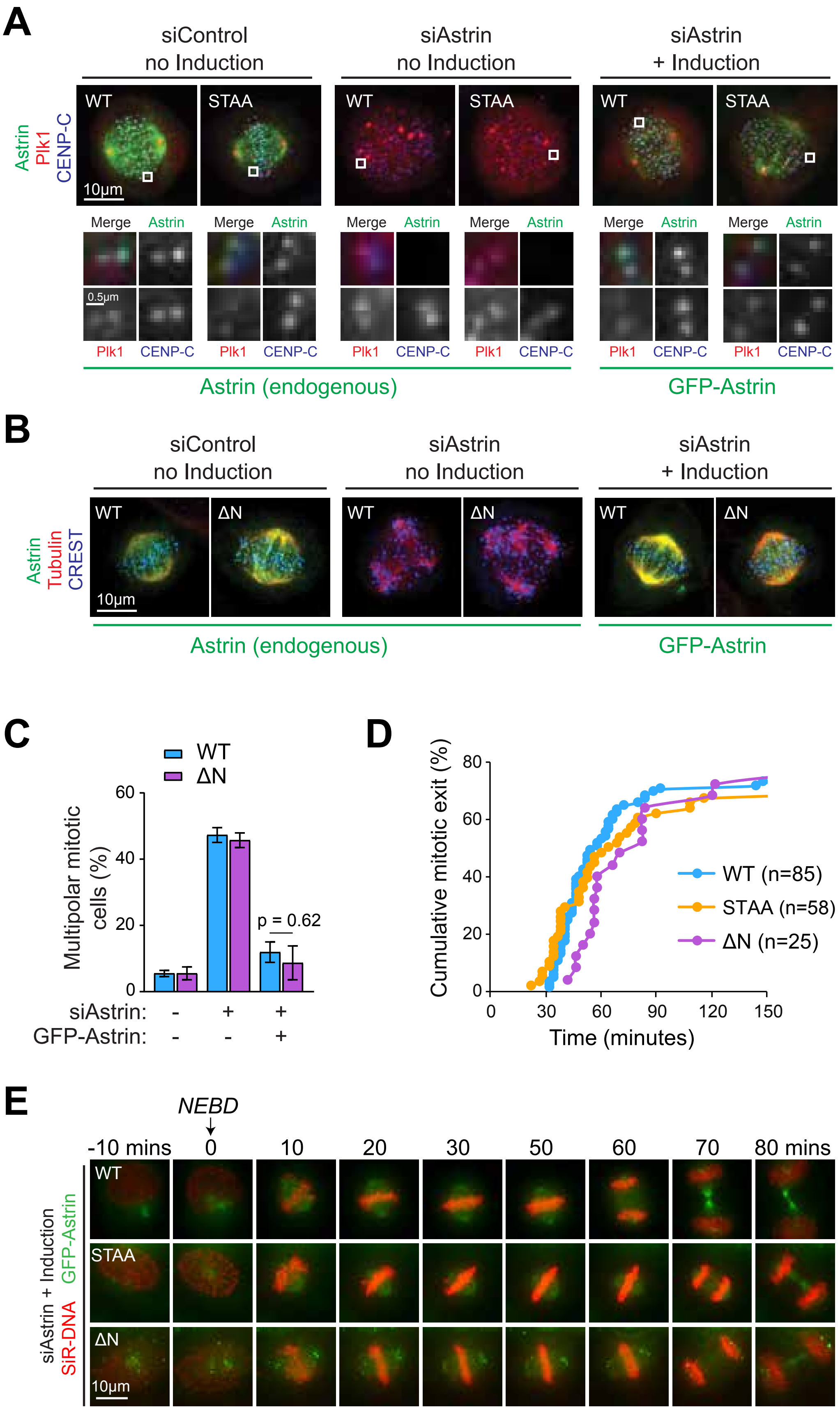
A
siControl no Induction
siAstrin no Induction
siAstrin + Induction
Astrin Plk1 CENP-C
WT
STAA
10µm
Merge Astrin
0.5µm
Plk1 CENP-C
Astrin (endogenous)
GFP-Astrin
B
siControl no Induction
siAstrin no Induction
siAstrin + Induction
Astrin Tubulin CREST
WT
ΔN
10µm
Astrin (endogenous)
GFP-Astrin
C
WT
ΔN
Multipolar mitotic cells (%)
0
20
40
60
p = 0.62
siAstrin: - + +
GFP-Astrin: - - +
D
Cumulative mitotic exit (%)
0
20
40
60
80
WT (n=85)
STAA (n=58)
ΔN (n=25)
0 30 60 90 120 150
Time (minutes)
E
NEBD
-10 mins 0 10 20 30 50 60 70 80 mins
WT
STAA
ΔN
siAstrin + Induction
GFP-Astrin
SiR-DNA
10µm

### Figure S4

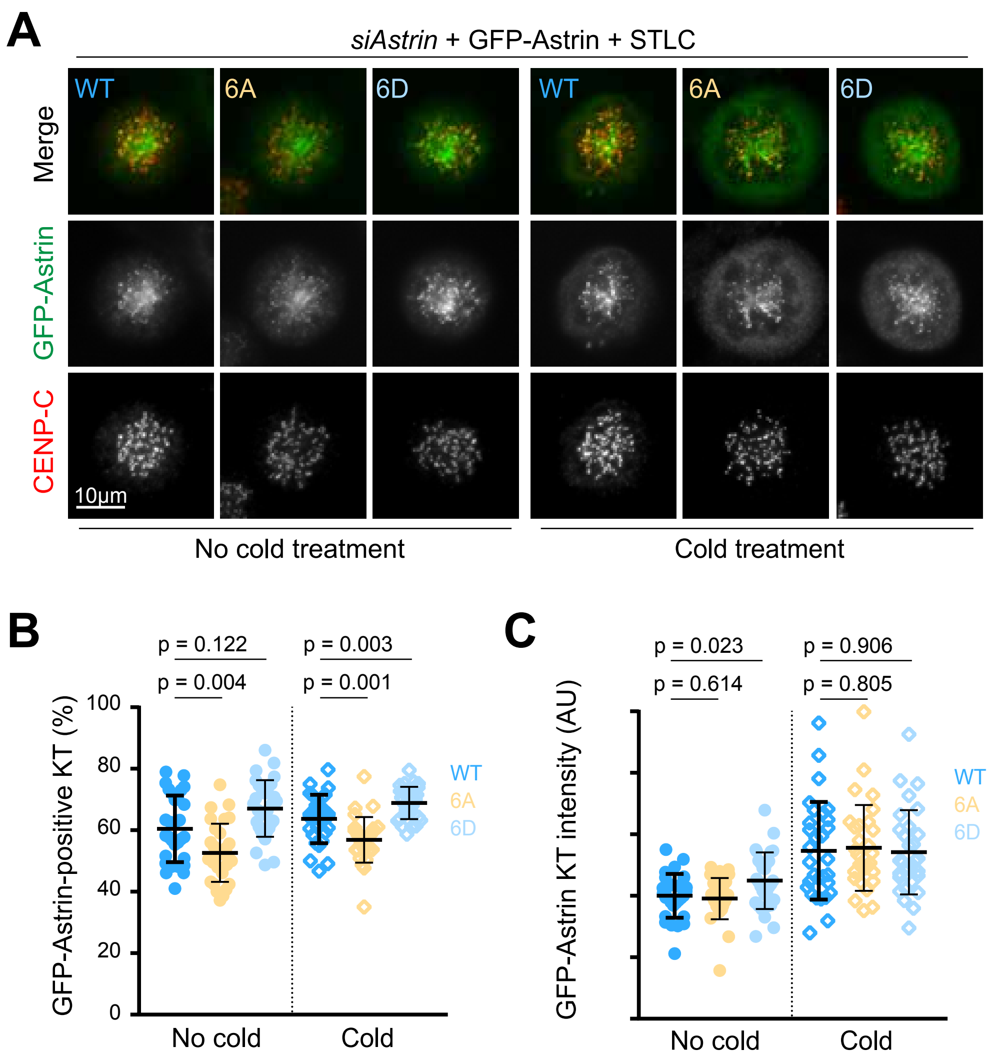
A
siAstrin + GFP-Astrin + STLC
WT
6A
6D
WT
6A
6D
Merge
GFP-Astrin
CENP-C
10µm
No cold treatment
Cold treatment
B
p = 0.122
p = 0.004
p = 0.003
p = 0.001
GFP-Astrin-positive KT (%)
100
80
60
40
20
0
WT
6A
6D
No cold
Cold
C
p = 0.023
p = 0.614
p = 0.906
p = 0.805
GFP-Astrin KT intensity (AU)
WT
6A
6D
No cold
Cold
